# Common-specific edge-centric connectome across Four Episodes in Bipolar Disorder

**DOI:** 10.1101/2025.10.03.680275

**Authors:** Xiaobo Liu, Sanwang Wang, Qiuxuan Yu, Xi-Han Zhang, Lang Liu, Yujun Gao

## Abstract

**Background:** Bipolar disorder (BD) is a heterogeneous psychiatric illness marked by dynamic mood states, including manic (BipM), depressive (BipD), mixed (mBD), and euthymic/remitted (rBD) episodes. These clinical fluctuations are accompanied by widespread functional disruptions in the brain. However, the shared and individual-specific neural mechanisms across distinct episodes of BD remain poorly understood.

**Methods:** We analyzed resting-state fMRI data from 190 participants (BD patients in four episodes and healthy controls) using edge-centric functional connectomes (eFC), which capture time-resolved co-fluctuations between brain regions. A common-orthogonal-basis extraction (COBE) algorithm was applied to decompose individual eFC matrices into shared and individual-specific subspaces. We characterized the spatial topology, genetic relevance, and circuit-level correlates of the shared component. Dynamic properties (entropy, identifiability) and symptom prediction models were assessed using entropy metrics, intraclass correlation coefficients, and support vector regression.

**Results:** The shared eFC pattern was stable across participants, aligned with the sensory–association gradient (*p* < 0.0001), and exhibited significant heritability and test–retest reliability (*p* < 0.05). Entropy of individual loadings increased with illness duration and was significantly elevated in BD, particularly in mBD. Microcircuit modeling revealed that this shared pattern was inversely related to external input strength (*r* = – 0.34, *p_spin_*< 0.05), indicating intrinsic network dominance. mBD was associated with globally elevated eFC entropy and markedly reduced fingerprint stability. Symptom severity (HDRS, YMRS, HAMA) was significantly predicted from individual network topographies across BD phases, highlighting clinically meaningful dynamic signatures.

**Conclusion:** Our findings demonstrate that BD episodes are underpinned by a conserved functional scaffold and distinct individual-specific neural fingerprints. Edge-centric dynamics—especially those derived from individual-specific decompositions—offer robust biomarkers for mood state characterization and symptom severity, and may facilitate future personalized interventions in BD.

## Introduction

Bipolar disorder (BD) is characterized by pronounced mood fluctuations, manifesting as manic (BipM), depressive (BipD), mixed (mBD) and euthymic/remitted (rBD) episodes (Grande et al., 2016a; Oliva et al., 2025; Pomarol-Clotet et al., 2015). These dynamic clinical states, such as affective instability, are frequently associated with a range of cognitive and behavioural impairments, as well as alterations in brain function (Liu, Wan, Zhang, Cui, et al., 2025a). Across individuals, BD is highly heterogeneous, exhibiting substantial variability in symptom profiles, neural activity, structural brain organization, and genetic architecture (Liu, Wan, Zhang, Cui, et al., 2025b; Liu, Wan, Zhang, Liu, et al., 2025). This inter-individual variability presents a major challenge for identifying consistent neurobiological mechanisms that distinguish among different BD phases.

Functional brain alterations in neuropsychiatric disorders are primarily characterized using functional MRI (fMRI) (Liu, Wan, Zhang, Cui, et al., 2025b, 2025c; Sommerfeldt et al., 2016; Y. Yang et al., 2021). Traditional “node-centric” connectomics infer brain organisation based on the synchrony of blood-oxygen-level–dependent (BOLD) signals across predefined brain regions (Betzel & Bassett, 2017; Bullmore & Sporns, 2009; Rubinov & Sporns, 2010). Recent methodological advances have introduced the *edge-centric functional connectome (eFC)*, which computes a time-resolved Pearson-correlation series for each pair of regions (Faskowitz et al., 2020; B. Yang et al., 2023). This approach captures dynamic fluctuations in connection strength, thereby providing a more direct and temporally nuanced readout of the brain’s communication dynamics. eFC analyses have already demonstrated enhanced sensitivity in identifying aberrant coordination in neuropsychiatric disorders such as autism (Anderson et al., 2011) and schizophrenia (Faskowitz et al., 2020; Herskovits et al., 2015; B. Yang et al., 2023), suggesting that eFC may uncover both dysregulated circuitry and biologically tractable therapeutic targets across the different phases of BD.

Traditional case–control designs emphasize group-level effects, implicitly assuming the existence of an “average patient” and, in doing so, often overlook the biological diversity and the phase-dependent heterogeneity inherent in BD. We propose that neural representations in BD comprise both (i) shared motifs and (ii) disease-specific heterogeneity. The human cortex conforms to a canonical sensory–association hierarchy that likely provides a macroscopic scaffold shared across both healthy individuals and those with BD (Liu, Wan, Zhang, Cui, et al., 2025b; Park et al., 2024). Superimposed upon this shared architecture are symptom-specific, heritable deviations in functional connectivity, giving rise to individualised dynamic fingerprints (Finn et al., 2015). These fingerprints have the potential to track illness progression (Pradhan et al., 2022) and demonstrate high heritability (Fotiadis et al., 2024; Pourmotabbed et al., 2024). Notably, functional-connectivity fingerprints have been shown to remain stable across the lifespan in large cohorts (St-Onge et al., 2023), with edge-centric approaches outperforming node-centric ones—underscoring the precision and individuality of topological organisation (Faskowitz et al., 2020; Troisi Lopez et al., 2025; B. Yang et al., 2023).

These findings suggest that neural representations across BD episodes can be conceptualized as a dual mapping: a shared network backbone that underpins affect regulation and cognitive control (Sporns, 2011, 2013b, 2013a), overlaid with fine-grained, microstructural (Sporns, 2013b) and genetic variation that modulates edge-wise signal amplitude and frequency. This interaction produces similar macroscopic imbalances across patients while preserving highly individualised edge-state perturbations.

To test this framework, we analysed resting-state fMRI from all four BD episodes using the *common-orthogonal-basis extraction* (COBE) algorithm (Shan et al., 2024). COBE decomposes multiple data blocks (in this case, eFC matrices) into a *common subspace* and *block-specific subspaces* (Zhou et al., 2016). For the common subspace and its loadings, we (i) examined their spatial correspondence with functional gradients, (ii) estimated heritability, and (iii) related the loadings to BD phase and clinical course. We further investigated how this common motif is linked to micro-circuit parameters. Finally, we assessed the fingerprinting capability of the individual-specific subspace and quantified the association between eFC strength within this subspace and clinical symptomatology through regression analyses.

## Results

### Decomposition of edge-centric functional connectome across BD subtypes

To address the heterogeneity of BD episodes and identify a shared neural communication structure, we decomposed eFC using the COBE method. This analysis builds upon prior parcellation and preprocessing of resting-state fMRI time series. COBE separated each subject’s eFC into shared and individual-specific components (Figure 1). The common pattern was consistently expressed across all BD subtypes—remitted (rBD, N = 37), depressive (BipD, N = 42), manic (BipM, N = 38), mixed (mBD, N = 38)—as well as healthy controls (HC, N = 35), enabling the dissociation of a group-level connectivity scaffold from personalized network dynamics. The eFC time series provided detailed temporal co-fluctuation information, and COBE isolated a stable shared subspace along with individualized loadings that were later used to inform symptom prediction.

**Figure. 1.**
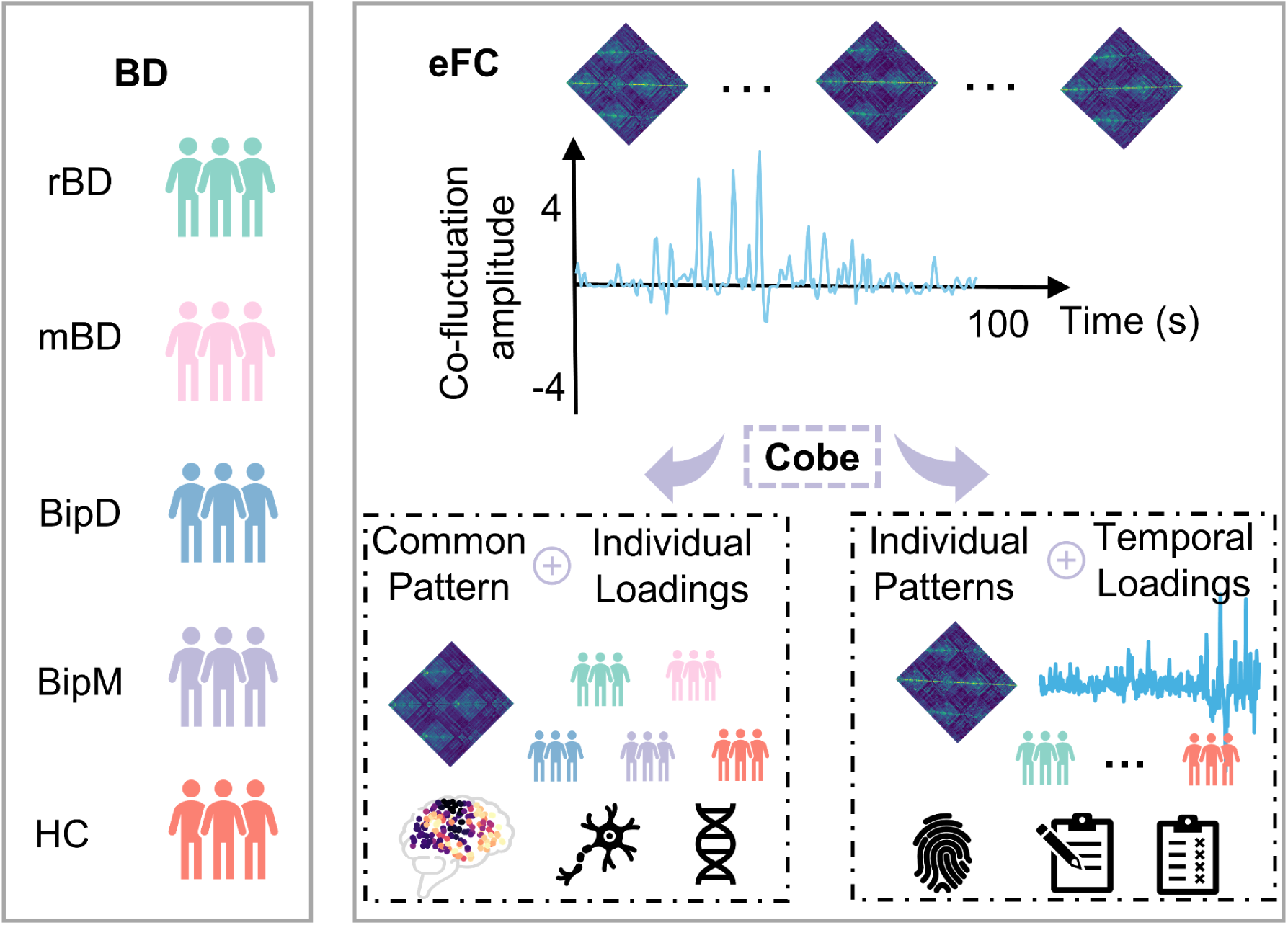
Schematic of the study workflow and COBE decomposition. Left: five cohorts comprising remitted (rBD, N = 37), mixed (mBD, N = 38), depressive (BipD, N = 42) and manic (BipM, N = 38) episodes of bipolar disorder, along with healthy controls (HC, N = 35). Right-top: edge-centric functional connectomes (eFC) derived from resting-state fMRI yield time-resolved co-fluctuation amplitude series for every pair of brain regions. Applying common-orthogonal-basis extraction (COBE) to the subject-specific eFC matrices separates (i) a common subspace with individual loadings, representing the shared communication scaffold, and (ii) individual-specific subspaces with temporal loadings, capturing personalised dynamic fingerprints that are linked to clinical symptoms.

### Spatial topology and genetic correlates of the shared eFC motif

To examine the neurobiological significance of the shared COBE-derived motif, we analyzed its spatial correlation with macroscale cortical functional gradients, evaluated its variability and heritability, and examined its association with regional gene expression profiles. This integrative analysis links large-scale functional topography to underlying genetic and developmental architecture.

The common COBE-derived component showed a significant positive correlation with cortical functional gradients (r = 0.41, p < 0.0001; Figure 2a), suggesting alignment with the established sensory-to-association hierarchy. To assess variability across groups, we quantified the Shannon entropy of individual component loadings (Figure 2b). Entropy was significantly elevated in all bipolar disorder (BD) subgroups compared to healthy controls (HC), reflecting increased dispersion and reduced stability of the shared pattern. Post hoc comparisons revealed distinct group-level differences: entropy in mBD was higher than in BipD (*t* = 4.92, *P_HSD_ <* 0.05) but lower than in HC (*t* = –9.84, *P_HSD_ <* 0.05); BipD showed greater entropy than BipM (*t* = 4.04, *P_HSD_ <* 0.05), HC (*t* = –4.51, *P_HSD_ <* 0.05), and rBD (*t* = 7.58, *P_HSD_ <* 0.05); BipM exhibited lower entropy than HC (*t* = –8.81, *P_HSD_ <* 0.05), but higher than rBD (*t* = 3.01, *p* < 0.05); and HC consistently exhibited greater entropy than rBD (*t* = 12.77, *P_HSD_ <* 0.05). Furthermore, entropy positively correlated with illness duration (r = 0.23, p = 0.0004; Figure 2c), suggesting a progressive destabilization of the shared motif. The common component also demonstrated significant heritability (H = 0.30, p < 0.05) and test-retest reliability (R = 0.51, p < 0.05), supporting its biological validity and temporal stability. Gene enrichment analyses revealed that spatial expression patterns of the common component was aligned with transcriptomic signatures related to synaptic signaling, ion transport, mitochondrial function, and neuronal system processes (Figure 2e).

**Figure 2.**
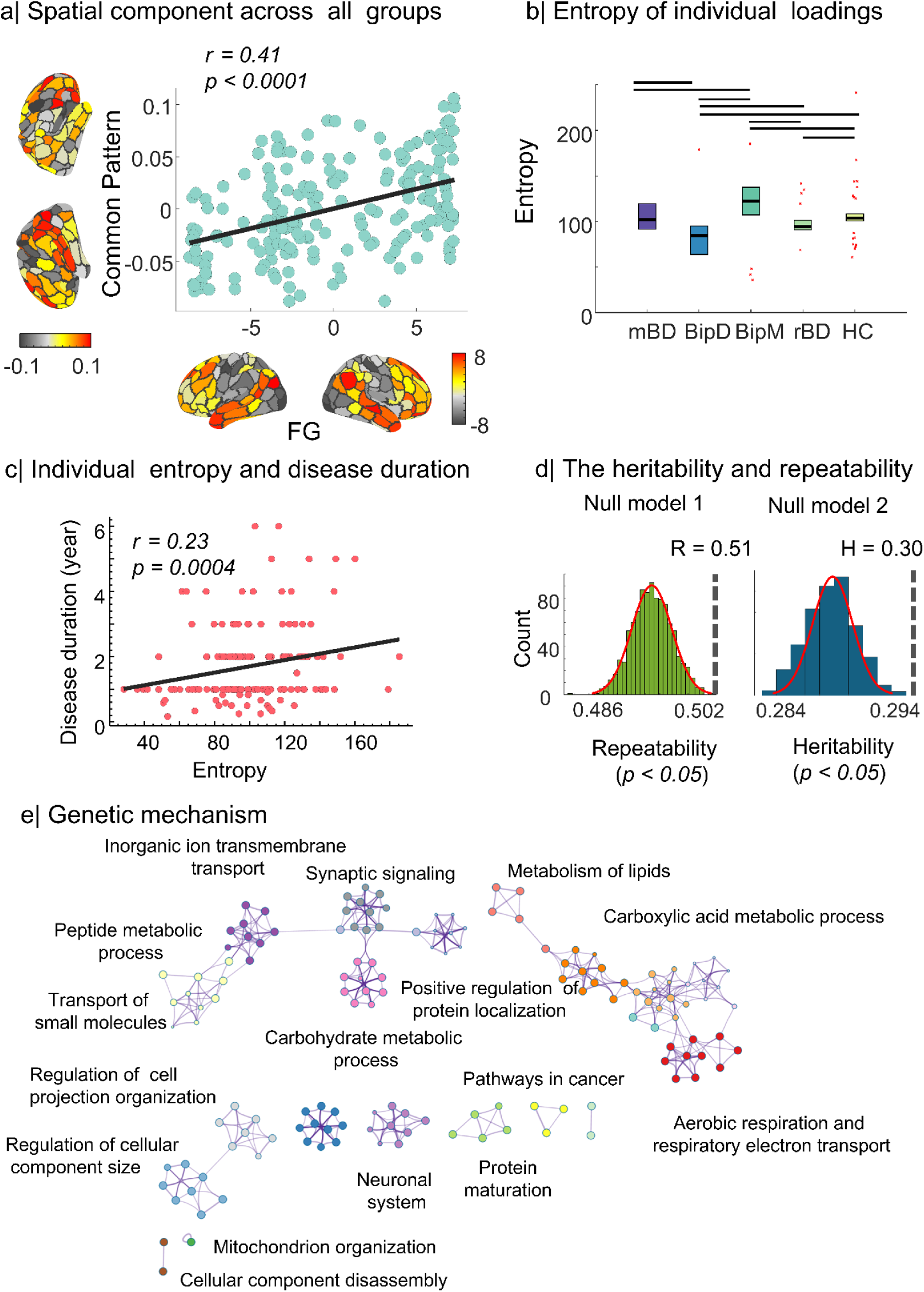
Functional topology, information-theoretic metrics, biological characteristics, and genetic basis of the shared connectivity component a. Spatial correspondence between the shared component and cortical functional gradients (FG): The cortical map depicts the cortical distribution of the common connectivity pattern identified via COBE across all groups. The scatterplot shows a significant positive correlation between the component loadings and functional gradients (*r* = 0.41, *p* < 0.0001), indicating that this shared motif follows the canonical sensory–association hierarchy. b. Entropy of individual loadings across diagnostic groups: Box plots display the Shannon entropy of individual loadings on the common component in different BD episodes(mBD, BipD, BipM, rBD) and healthy controls (HC). All BD subgroups exhibit significantly higher entropy than HC, reflecting increased variability and reduced stability in the expression of the shared motif. b. Association between individual entropy and illness duration: A positive correlation was observed between entropy of individual loadings and illness duration (*r* = 0.23, *p* = 0.0004), suggesting that longer disease duration may be associated with destabilization of the shared functional architecture. c. Repeatability and heritability of the shared component: The shared component demonstrated significant test–retest reliability (*R* = 0.51, *p* < 0.05) and heritability (*H* = 0.30, *p* < 0.05), indicating that this spatial motif is both stable across measurements and partly genetically determined. d. Gene ontology enrichment of the common component: Network graph displays biological processes enriched in the genetic profile associated with the common connectivity pattern, including synaptic signaling, ion transmembrane transport, metabolic processes (e.g., lipids, peptides, carbohydrates), cellular organization, mitochondrial function, and neuronal development—highlighting the neurobiological relevance of this shared functional scaffold.

### Relationship between common connectivity and circuit-level mechanisms

To examine whether the shared functional motif reflects underlying circuit properties, we simulated neural dynamics using a biophysically constrained excitatory-inhibitory (E–I) recurrent network model, informed by group-level structural connectivity (Figure 3, top). From the model, we derived regional parameters indicative of recurrent connection strength (w) and external input (I) (Figure 3, middle).

**Figure 3.**
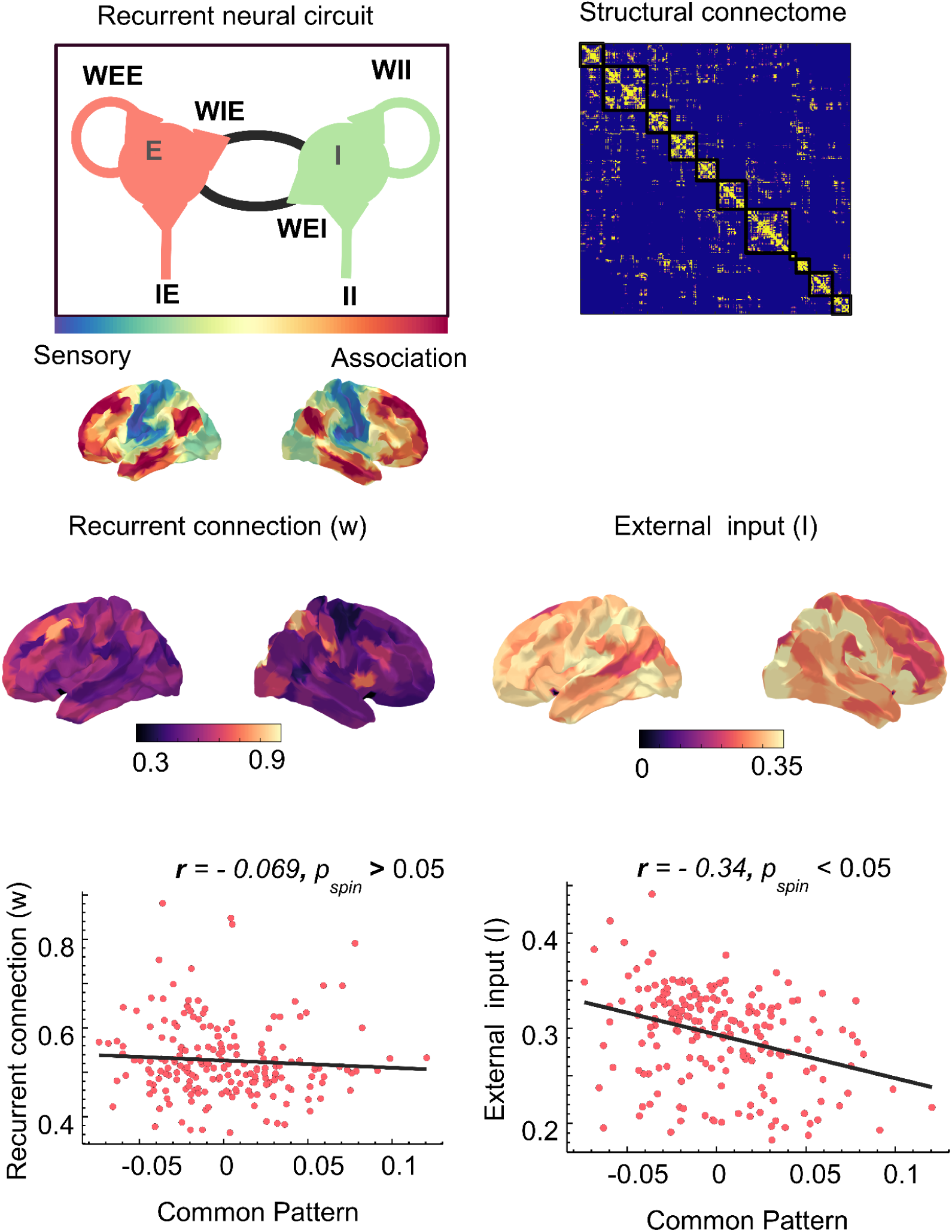
Association between the shared functional motif and microcircuit parameters derived from a recurrent neural model Top-left: Schematic of a recurrent excitatory–inhibitory (E–I) circuit model incorporating intra-population (WEE, WII), inter-population (WIE, WEI) connections, along with external input parameters(IE, II), used to estimate region-specific circuit characteristics. Top-right: Group-averaged structural connectome used to constrain model simulations. Middle row: Cortical surface projections of two key model-derived parameters: recurrent connection strength (w) and external input (I). The sensory–association axis is overlaid to illustrate the hierarchical topography along which these parameters are distributed. Bottom-left: Scatterplot showing no significant spatial association between recurrent connection strength and expression of the common functional pattern (*r* = –0.069, *p_spin_* > 0.05). Bottom-right: Scatterplot showing a significant negative spatial correlation between external input and the shared functional motif (*r* = –0.34, *p_spin_* < 0.05), suggesting that regions strongly involved in the shared network scaffold may rely more on intrinsic dynamics and less on exogenous drive.

Spatial correspondence analyses revealed that recurrent excitation was not significantly associated with the common component (*r* = –0.069, *p_spin_* > 0.05). In contrast, external input strength showed a significantly negative correlation with the common motif (*r* = –0.34, *p_spin_* < 0.05; Figure 3, bottom). These findings suggest that regions with stronger expression of the shared functional pattern may rely less on exogenous input and more on intrinsic network dynamics.

### Dynamic instability and identifiability of edge-centric fingerprints across BD subtypes

To evaluate the differences in eFC temporal dynamics across clinical states in BD, we quantified the entropy of edge-wise co-fluctuations and calculated fingerprint reliability using ICC. This expands upon the individual-level decompositions and assesses temporal and diagnostic variations.

Edge-wise entropy matrices indicated increased dynamic variability in the mBD group compared to other BD subtypes and HC (Figure 4a). ICC analysis demonstrated that fingerprint stability was highest in HC and progressively decreased across rBD, BipD (*t =* 3.31, *P_HSD_ <* 0.05), BipM (*t =* 3.30, *P_HSD_ <* 0.05), and was lowest in mBD (*t = 4.14*, *P_HSD_ <* 0.05) (Figure 4b). Furthermore, radar plots of entropy across canonical resting-state networks showed disproportionately elevated entropy in mBD across all networks, particularly within frontoparietal and salience systems (Figure 4c), indicating a widespread instability of edge dynamics during this period.

**Figure 4.**
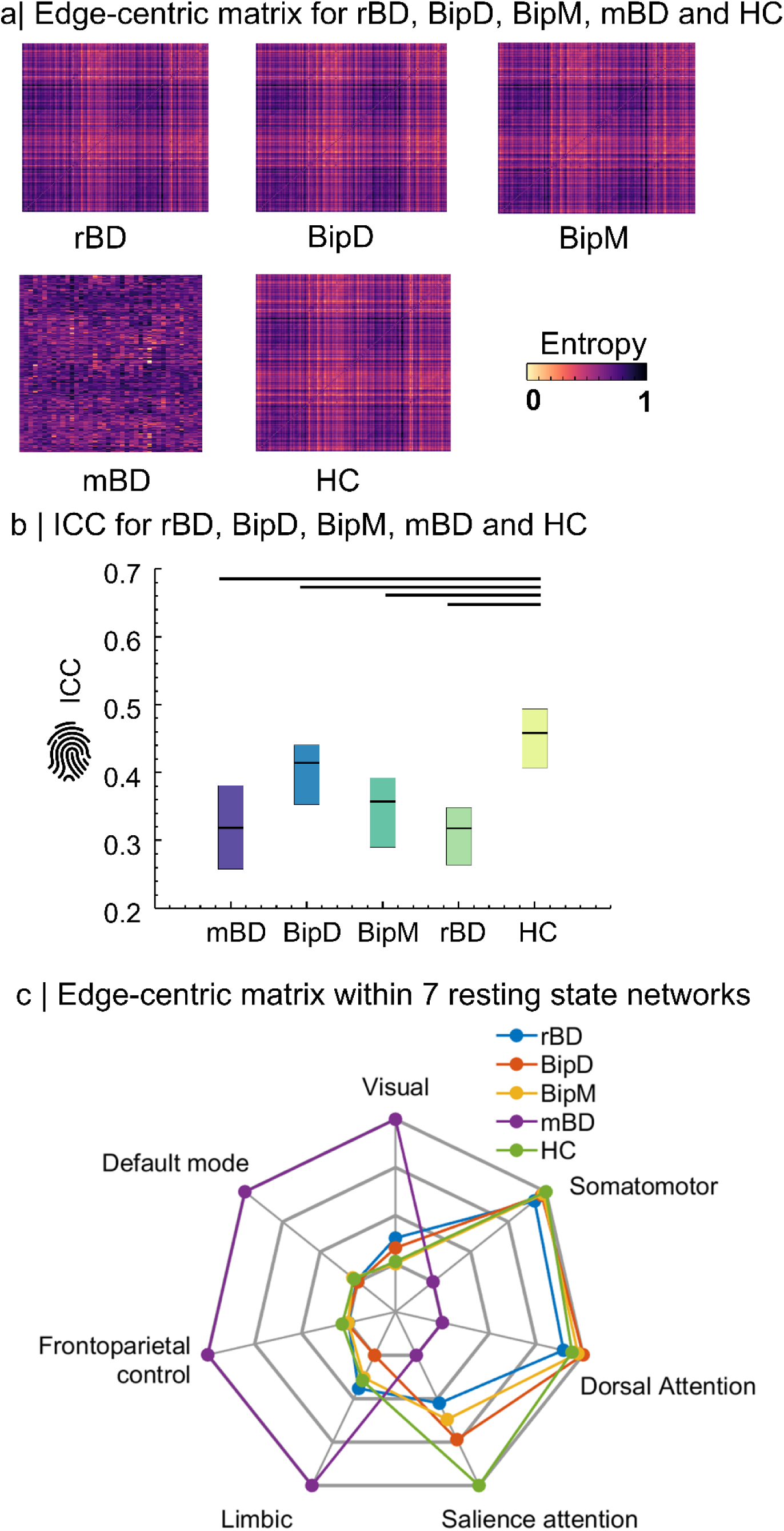
Edge-centric connectome entropy and fingerprint stability across BD episodes and healthy controls a. Group-averaged edge-centric entropy matrices Edge-centric entropy matrices represent the temporal variability (entropy) of edge-wise co-fluctuations for each diagnostic group: rBD, BipD, BipM, mBD, and HC. The mBD group demonstrates prominently elevated entropy, indicating greater dynamic instability in functional coordination compared to other BD subtypes and HC. b. ICC of edge-centric fingerprints Box plots show the intraclass ICC of edge-centric connectivity profiles within each group, reflecting the reliability of edge-centric fingerprints across sessions. HC and rBD show the highest identifiability, whereas mBD displays the lowest, suggesting impaired stability of functional signature during active mixed episodes. Horizontal lines indicate statistically significant pairwise differences (*p* < 0.05). c. Network-specific entropy profiles across seven canonical resting-state networks Radar plots display the average edge-centric entropy across seven canonical resting-state networks: visual, somatomotor, dorsal attention, salience, limbic, frontoparietal control, and default mode networks. Compared to other groups, mBD demonstrates consistently elevated entropy across all networks—particularly in higher-order control systems—indicating widespread dysregulation and instability of large-scale communication dynamics during mixed states.

### Clinical relevance of individual-specific network topographies

To assess whether individualized network dynamics predict symptom load, we trained machine learning models—specifically support vector regression (SVR) and random forest regression—utilizing personalized COBE features to estimate HAMA, HDRS, and YMRS scores. This analysis builds on the individual-specific components and correlates them with clinical phenotypes.

Multivariate SVR models trained on individualized eFC patterns significantly predicted HAMA scores in rBD (*r* = 0.40, *p* = 0.015), HDRS scores in BipD (*r* = 0.35, *p* = 0.023) and mBD (*r* = 0.34, *p* = 0.038), and YMRS scores in BipM (*r* = 0.38, *p* = 0.020) (Figure 5). These findings indicate that individualized eFC-derived topographies encapsulate clinically relevant variance and may serve as potential biomarkers for stratifying symptom burden across BD phases.

**Figure 5.**
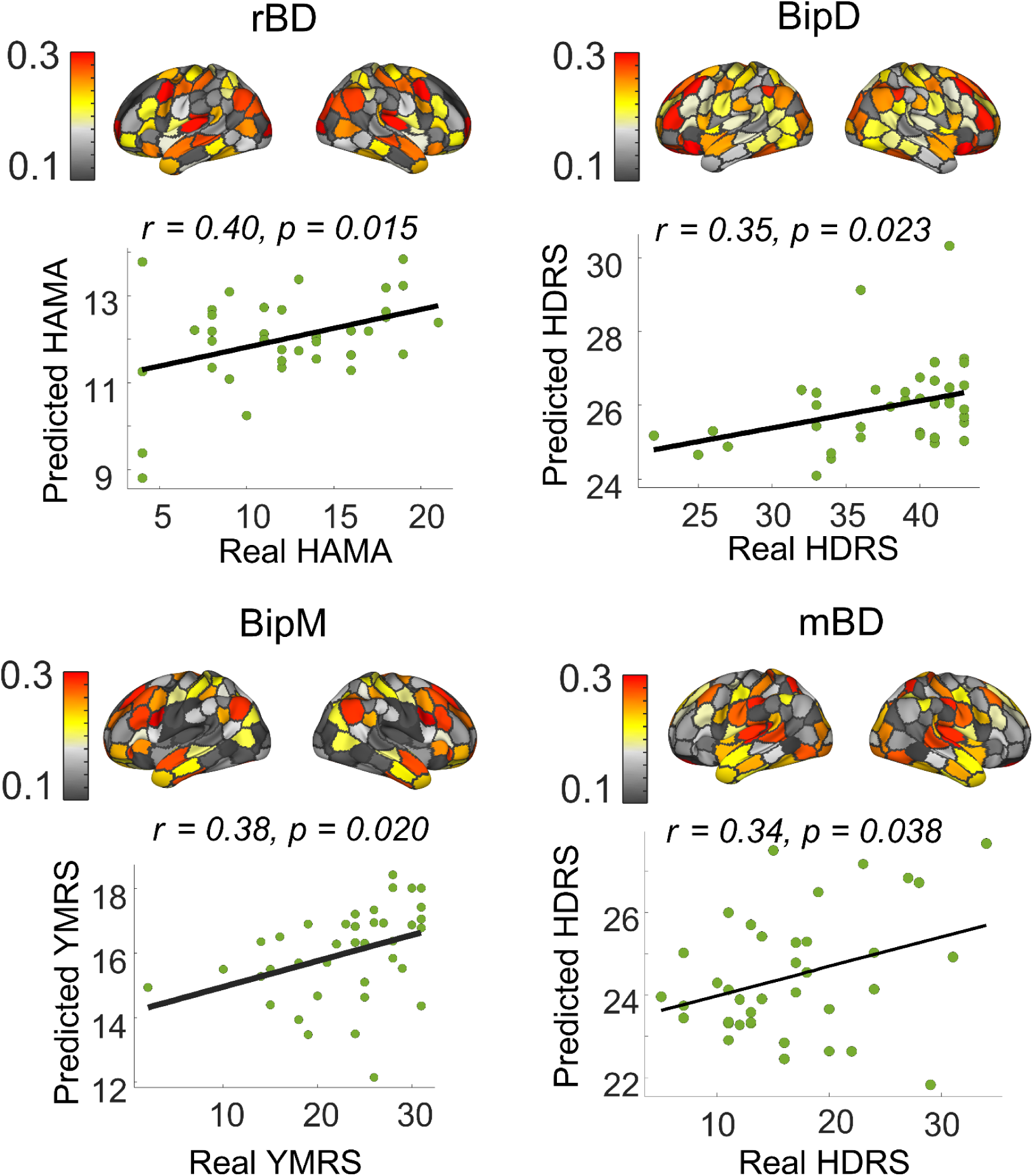
Associations between individual-specific network topographies and clinical symptom severity across BD phases Spatial distributions of edge-centric functional connectivity features derived from individualized COBE subspaces are shown for each BD phase—remitted (rBD), depressive (BipD), manic (BipM), and mixed (mBD)—is illustrated in the top row, with symptom prediction performance depicted in the bottom row. Brain maps illustrate the strength of edge-centric functional connectivity features obtained from individual-specific COBE subspaces. rBD (top-left): Anxiety severity (HAMA) was significantly predicted from individualised network patterns (*r* = 0.40, *p* = 0.015), suggesting residual anxiety-related network signatures in the euthymic state. BipD (top-right): Depression severity (HDRS) was predicted from individual network topology (*r* = 0.35, *p* = 0.023), highlighting phase-specific neural alterations associated with depressive symptoms. BipM (bottom-left): Manic symptom severity (YMRS) showed a significant association with individualised network configurations (*r* = 0.38, *p* = 0.020), indicating their relevance to mania-related dysfunction. mBD (bottom-right): HDRS scores were predicted from network topology in the mixed state (*r* = 0.34, *p* = 0.038), demonstrating symptom-specific neural patterns even in complex, affectively unstable episodes.

## Discussion

This study systematically investigated the shared and individual-specific neural mechanisms underlying dynamic brain function across different episodes of BD—including BipM, BipD, mBD, and rBD episodes—by leveraging eFC and the COBE framework. Our findings indicate a dual-axis architecture of functional brain organization in BD: (i) a conserved, cross-individual communication scaffold that reflects a stable functional core across disease phases, and (ii) phase-specific, individualized dynamic variations directly linked to the severity of clinical symptoms. This framework advances our understanding of the macroscale neurobiology of BD by disentangling stable and flexible dimensions of brain network architecture, offering a biologically grounded basis for stratified diagnosis and personalized treatment approaches.

### Shared Functional Structure Revealed by eFC Decomposition: Stability, Biological Substrate, and Topographical Alignment

Using the COBE framework on high-dimensional eFC data, we firstly identified a shared functional connectivity pattern consistently expressed across all BD subtypes and healthy controls. Spatially, this pattern exhibited strong alignment with the canonical sensory–association gradient, recapitulating the hierarchical macroscale organization of the cortex. This finding broadens the application of the functional gradient (Dong et al., 2021; Margulies et al., 2016) and indicates that such a hierarchical structure remains preserved even in the context of severe psychiatric illness.

Additionally, the shared pattern demonstrated strong consistency and notable heritability, suggesting that it represents a stable macroscale scaffold that is both trait-like and genetically constrained. This aligns with the “functional fingerprint” hypothesis by Finn et al. (2015), which posits that functional connectivity patterns are both highly individualized and stable across time. Gene enrichment analyses further revealed that the spatial topography of this shared pattern was significantly associated with transcriptional signatures involved in synaptic signaling, mitochondrial function, and neurodevelopmental processes, highlighting the biological validity of this conserved scaffold.

### Entropy as an Index of Disease Progression and Dysregulated Functional Modulation

The entropy of individual expression of the shared functional motif was significantly elevated in BD patients and positively associated with illness duration, suggesting progressive instability in the recruitment of this large-scale communication scaffold over the course of disease. These findings are consistent with the neuroprogression hypothesis (Berk et al., 2011), which posits that repeated mood episodes and chronicity in BD lead to cumulative disruptions in brain function and structure.

To our knowledge, this is the first study to employ Shannon entropy as a metric for quantifying the flexibility and variability of engagement with a shared functional network, thereby extending beyond traditional static measures such as variance or temporal fluctuations. Elevated entropy may represent a maladaptive shift in the balance between flexibility and integration, indicative of a noisy and inefficient communication regime. In this context, entropy captures a core aspect of dynamic dysfunction and may offer a quantifiable biomarker for tracking illness progression and evaluating treatment response.

### Circuit-Level Mechanism of the Shared Component: Reduced External Input Suggests Intrinsic Regulation

To further elucidate the physiological basis of the shared motif, we employed a biophysically informed recurrent excitatory–inhibitory (E–I) neural circuit model. This modeling approach revealed a significant negative association between the expression of the shared functional component and the strength of external input, while no significant relationship was observed with recurrent excitation. These results suggest that the common connectivity scaffold is more likely governed by intrinsic cortical dynamics rather than exogenous afferent drive.

This observation supports the “self-organizing brain” hypothesis (Deco et al., 2013; Honey et al., 2009), which suggests that large-scale resting-state connectivity patterns emerge from endogenous neural activity. AccordinglyIn this context, the preserved shared scaffold observed across all BD phases and healthy controls may represent a genetically and developmentally constrained functional baseline that supports core processes such as affective regulation and cognitive control. Disruption of this intrinsically maintained architecture may thus contribute to the emergence of transdiagnostic psychopathology.

### Mixed Episodes Exhibit Profound Network Instability and Fingerprint Disintegration

Entropy analyses revealed that the mixed episode (mBD) group exhibited the highest edge-level functional instability and the lowest fingerprint identifiability (intraclass correlation coefficient, ICC) across all groups. These findings indicate substantial disruption in dynamic coordination and a disintegration of individual-specific functional architecture. The instability was particularly prominent within the frontoparietal control, salience, and default mode networks—core systems responsible for cognitive integration and affective regulation.

These results provide strong empirical support for the “disintegration hypothesis” for mBD (Phillips & Swartz, 2014), which posits that mBD reflects a collapse of network-level coordination. Notably, our study fills a critical gap in the neurobiological characterization of mBD, a clinically severe yet understudied BD subtype. The pronounced dynamic disruption observed here not only highlights candidate neural circuits for targeted intervention but also offers a potential network-level biomarker to aid in the differential diagnosis and clinical management of mixed episodes.

### Individualized Dynamic Connectivity Predicts Multidimensional Symptomatology

Using support vector regression (SVR), we demonstrated that individual-specific connectivity patterns extracted from COBE-derived residual subspaces reliably predicted symptom severity across multiple clinical dimensions—namely, anxiety (HAMA) in rBD, depression (HDRS) in BipD and mBD, and mania (YMRS) in BipM. These findings suggest that fine-grained edge-level dynamics fluctuations encode clinically relevant information specific to each phase of BD.

Compared to prior studies employing static node-based functional connectivity (e.g., Drysdale et al., 2017), our time-resolved edge-centric framework captures more nuanced neural signals linked to transient symptomatic states. The ability to predict dimensional symptom severity from individualized neural fingerprints supports the translational utility of this approach. Specifically, dynamic eFC-based models offer a promising avenue for phase-specific identification, risk stratification, and prognosis modeling in BD and broader psychiatric populations.

## Limitation

This study has several limitations. First, although all four major episodes of bipolar disorder were included, the overall sample size was relatively modest, which may limit statistical power and reduce the generalizability of subgroup-specific findings. Future research should aim to recruit larger, demographically diverse cohorts and adopt longitudinal designs to better capture dynamic changes over time. Second, while the eFC approach enables the detection of subsecond fluctuations in functional connectivity, it remains constrained by the temporal resolution of fMRI and cannot capture faster neural dynamics (e.g., gamma-band oscillations activity). Integration with modalities such as MEG or EEG would help overcome this limitation and enable a more comprehensive assessment of neural dynamics. Third, the COBE algorithm operates under linear assumptions and may not fully model the complex, non-linear interactions that characterize brain networks. Future work could explore non-linear dimensionality reduction techniques, such as variational autoencoders or manifold learning, to improve the modeling of individualized network dynamics. Fourth, although spatial associations between the shared connectivity pattern and gene expression were identified, these findings remain correlational. Causal or longitudinal links with disease progression have yet to be established, underscoring the need for mechanistic and interventional follow-up studies. Finally, while the predictive models demonstrated statistical significant associations with symptom severity, their clinical utility and robustness have not been validated in external and multi-site datasets. Future efforts should prioritize the development of generalizable, clinically reliable models for deployment in real-world psychiatric settings.

## Conclusion

We propose and empirically validate a dual-representation model of functional brain organization in BD: a conserved, genetically influenced communication scaffold overlaid with individualized, state-dependent edge-level dynamic perturbations. This framework reconciles the long-standing paradox of reproducible macroscopic disruptions coexisting with pronounced individual variability in psychiatric illness. Our findings highlight the utility of edge-centric dynamic connectomics—especially when combined with COBE decomposition—as a scalable and biologically grounded tool for personalized symptom profiling in BD. Importantly, this approach offers a bridge between population-level neuroimaging and individualized clinical neuroscience. Future studies should prioritize: (1) longitudinal monitoring of edge dynamics to predict phase transitions or relapse risk; (2) integration of multimodal data (e.g., genomics, metabolomics, neurochemical imaging) to delineate mechanistic pathways; and (3) development of real-time, closed-loop neuromodulation protocols (e.g., fMRI neurofeedback or TMS) guided by individualized edge-state biomarkers. In summary, the eFC–COBE framework provides a novel, dynamic lens to map the neural landscape of BD and bridges the gap between population-based neuroimaging and individualized psychiatric care—marking a critical advance from “average brain mapping” to “dynamically individualized neurophenotyping” in precision psychiatry.

## Method

This study was approved by the Ethics Committee of Renmin Hospital of Wuhan University (Approval No. WDRY22022-K195). Prior to enrollment, detailed information regarding the study’s objectives, procedures, and potential risks was provided to all participants and their legal guardians, particularly in cases involving individuals with limited decision-making capacity. Informed consent was obtained only after ensuring that participants and their families had a comprehensive understanding of the study. Written consent was subsequently documented. The study was registered with both the National Medical Research Registration Information System and the Chinese Clinical Trial Registry (ChiCTR2200064938).

### Clinical symptoms and cognitive assessment

The Young Mania Rating Scale (YMRS) is a clinician-rated instrument developed to quantify the severity of manic symptoms. It comprises 11 items, with items 1, 2, 3, 4, 7, 10, and 11 rated on a 5-point scale (0–4), and items 5, 6, 8, and 9 rated on a 9-point scale (0–8), allowing for differential weighting of symptom dimensions (Young et al., 1978). Each YMRS item is rated by clinicians based on patient interviews and observed behavior, with the total score providing a standardized index of manic symptom severity to inform clinical evaluation and treatment planning. Anxiety symptoms were independently assessed using the Hamilton Anxiety Rating Scale (HAMA), a validated instrument for the objective quantification of anxiety severity. The HAMA consists of 14 items, each rated on a 5-point scale ranging from 0 (not present) to 4 (very severe), reflecting the intensity of anxiety symptoms. The scale captures two primary dimensions of anxiety—psychic (mental agitation and psychological distress) and somatic (physical complaints related to anxiety)—which are analyzed separately to delineate symptom profiles (Jiang et al., 2025). The Hamilton Depression Rating Scale (HDRS) is a widely utilized clinician-administered measure for evaluating the severity of depressive symptoms (Hamilton, 1960). The 17-item version (HDRS-17) is commonly employed to classify depression severity, with lower total scores indicating milder symptomatology and higher scores reflecting more severe depressive states. The HDRS can be parsed into seven symptom clusters: Anxiety/Somatization, Weight, Cognitive Disturbance, Diurnal Variation, Psychomotor Retardation, Sleep Disturbance, and Feelings of Despair. Functional impairment was assessed using the Perceived Disability Questionnaire (PDQ), a 20-item self-report measure designed to evaluate perceived difficulties in daily functioning, including occupational, social, and leisure activities. Each item is rated on a 5-point Likert scale (0 = no difficulty; 4 = extreme difficulty), capturing functional domains such as concentration, task completion, interpersonal engagement, and routine responsibilities.

### Imaging Dataset

Data were collected from individuals diagnosed with BD who sought treatment at the Department of Psychiatry and Psychology, Renmin Hospital of Wuhan University, between September 2021 and December 2023. Diagnostic screening was conducted using the Chinese version of the Mini-International Neuropsychiatric Interview (MINI) (Amorim, 2000). Formal diagnoses were confirmed by two board-certified psychiatrists in accordance with DSM-5 criteria for BD (Association & others, 2000). Participants were stratified into clinical subgroups based on established criteria. For the manic bipolar disorder group (Grande et al., 2016b): age 18–45 years, DSM-5 diagnosis of BD, YMRS score >8, HDRS-17 score <7, and either first-episode untreated or undergoing treatment for the first time. The depressed bipolar disorder group (Tohen et al., 2012) included patients with HDRS-17 >12 and YMRS <7. The rBD group was defined by age 18–45, HDRS-17 <7, and YMRS <7. The mBD group met the following criteria: age 18–45, DSM-5 diagnosis of BD, HDRS-17 >12, YMRS >8, and self-initiated medication discontinuation exceeding 14 days. **Exclusion criteria** for all BD subgroups included contraindications to MRI, history of organic brain disease, comorbid psychiatric diagnoses, history of pharmacological or physical treatment (for BipM only), left-handedness, unstable medical conditions, history of substance abuse, pregnancy or lactation, and concurrent neurological disorders. Age- and sex-matched HCs were recruited from the local community, universities, and via posters at Hubei Provincial People’s Hospital. All HCs were right-handed and screened to exclude individuals with MRI contraindications, neurological conditions, psychiatric history, substance abuse, pregnancy or lactation, or a family history of psychiatric or neurological disorders. **Withdrawal and termination criteria** were clearly defined. Participants could withdraw at any time by revoking informed consent. Investigators also retained the right to discontinue participation if subjects were deemed unsuitable to continue. Termination criteria included noncompliance resulting in invalid data, exclusion from final analyses, or inability to complete the study protocol.

Neuroimaging data were acquired using a 3T GE SIGNA Architect MRI scanner equipped with a 48-channel head coil. During scanning, participants were instructed to remain still, stay awake, relax, and keep their eyes closed. Foam padding and earplugs were used to minimize head motion (threshold was 2 mm) and acoustic discomfort. All scans were performed by two certified MRI technologists with intermediate-level professional qualifications. High-resolution T1-weighted anatomical images (T1_3D) were acquired using the following parameters: maximum repetition time (TR), minimum echo time (TE), number of excitations (NEX) = 1, slice thickness = 2 mm, and field of view (FOV) = 256 × 256 mm². Total scan duration was 7 minutes. Resting-state functional MRI (rs-fMRI) data were collected using an echo-planar imaging (EPI) sequence with the following parameters: TR = 2000 ms, TE = 30 ms, FOV = 220 × 220 mm², flip angle = 90°, matrix = 64 × 64, in-plane resolution = 3 × 3 × 3 mm³, slice thickness = 36 mm, and 240 time points. The rs-fMRI scan duration was 9 minutes. The demographics were shown in **S-Table 1**. And the quality control could be seen in S-Figure 1.

### Data preprocessing

For all datasets, raw DICOM files were converted to Brain Imaging Data Structure (BIDS) format using HeuDiConv v0.13.1. Structural and functional MRI preprocessing was performed using *fMRIPrep* version 23.0.2 (Esteban et al., 2019), a robust pipeline built upon Nipype v1.8.6 (Gorgolewski et al., 2011). Anatomical preprocessing included intensity non-uniformity correction, skull stripping, tissue segmentation, cortical surface reconstruction, and spatial normalization to MNI space. Functional preprocessing comprised motion correction, slice timing correction, susceptibility distortion correction (when applicable), co-registration to the corresponding T1-weighted image, and transformation to standard space. Full preprocessing reports were generated by *fMRIPrep* for quality assessment. Preprocessed time series were parcellated into 200 cortical regions based on the Schaefer 200×7 functional atlas. Subsequent nuisance regression was performed using *Nilearn*, following the “simple” confound removal strategy described by (Wang et al., 2024).

### Edge-centric functional connectome (eFC) establishment

Using the Schaefer-200 atlas, cortical areas were parcellated into 100 distinct regions, from which regional time series were extracted and subsequently z-scored. To characterize functional connectivity dynamics, we computed edge time series by calculating the dot product of the time series between each pair of these 200 nodes. This procedure yielded edge-specific time series for all 19,900 unique node pairs (200 × 199/2).

### Edge-centric functional connectome (eFC) establishment

Using the Schaefer-200 atlas, cortical areas were parcellated into 100 distinct regions, from which regional time series were extracted and subsequently z-scored. To characterize functional connectivity dynamics, we computed edge time series by calculating the dot product of the time series between each pair of these 200 nodes. This procedure yielded edge-specific time series for all 19,900 unique node pairs (200 × 199/2).

### Common orthogonal basis extraction (COBE) method

Furthermore, we applied the COBE method on the edge time series of the rBD, BipD, BipM, and HC groups to decomposition partitions the data into a **common subspace**, shared across all subjects, and **individual-specific subspaces**, capturing variability at the single-subject level. COBE identifies a set of orthogonal bases in which edge time series across all subjects can be efficiently represented, revealing connectivity patterns consistently expressed at the group level. This approach effectively captures shared features across datasets while simultaneously preserving individual-specific variations, enabling the separation of common connectivity patterns from unique individual characteristics for subsequent analyses.

### Stability and heritability of eFC

In this study, we evaluated *eFC* through two main aspects: test-retest reliability and heritability, which capture temporal stability and genetic contributions, respectively. Using the Human Connectome Project (HCP) S1200 data set (Elam et al., 2021), we analyzed rs-fMRI data from 1014 healthy adults (470 males, mean age 28.7 years), including 298 monozygotic and 188 dizygotic twins, alongside 720 singletons. We also included a subset of 45 retest subjects (13 males) scanned after a 140-day interval.

Test-retest reliability of *eFC* was assessed using the intraclass correlation coefficient (ICC) (Shrout & Fleiss, 1979), calculated as the ratio of intra-subject variance to total variance. We further examined the proportion of significant edges among total edge patterns post-FDR correction to interpret reliability. Heritability of *eFC* was estimated using SOLAR (v8.5.1b) (Almasy & Blangero, 1998), which models the covariance among family members based on genetic proximity. Narrow-sense heritability (*h²*) quantifies the proportion of phenotypic variance attributable to additive genetic factors. We incorporated age, sex, and age-related covariates into our models and averaged heritability across two sessions to generate final estimates.

### Enrichment Analysis for Common Spatial Patterns

To explore the molecular correlates of the common functional topography, we integrated spatially resolved gene expression data from the Allen Human Brain Atlas (AHBA; http://human.brain-map.org) (Shen et al., 2012). The dataset comprises 3,702 microarray samples from six adult postmortem donors (mean age = 42.5 ± 13.4 years; five males), sampled across anatomically distributed cortical regions. Gene expression data were processed using the *Abagen* toolbox (https://github.com/netneurolab/abagen) (Markello & Misic, 2021) and mapped to the Schaefer 400-region cortical parcellation.

To estimate cell-type-specific contributions, transcriptomic deconvolution was performed to infer the relative proportions of distinct cell types from bulk microarray data. Spatial correlations were then calculated between regional gene expression profiles and the common functional gradient to identify genes aligned with the spatial topography. To assess the biological relevance of these associations, we examined whether genes previously reported as transcriptionally dysregulated in postmortem cortical tissue were preferentially expressed in spatially aligned regions. Enrichment analyses for genes linked to functional reorganization were conducted using *Metascape* (Zhou et al., 2019), which integrates over 40 curated biological databases (https://metascape.org). Gene sets were thresholded at a false discovery rate (FDR)–corrected significance level of q < 0.05 and evaluated against a spatial null model to control for autocorrelation effects.

To further assess spatial correspondence between common functional gradients and neurobiological features, we employed a spatial permutation framework designed to preserve spatial autocorrelation while disrupting topographic alignment (Hansen et al., 2022b). Cortical receptor density maps were initially correlated with the common spatial pattern. A null distribution was generated by applying random spherical rotations to cortical maps, reassigning values to the nearest rotated parcels. To maintain anatomical symmetry, each rotation was applied to one hemisphere and mirrored to the contralateral side. This procedure was repeated 1,000 times. Empirical correlations were compared against the resulting null distribution, and statistical significance was determined as exceeding the 95th percentile of the null distribution derived from both spatial and temporal surrogates. This approach provides robust inference regarding topographic associations while explicitly accounting for spatial dependencies inherent in cortical data.

### Individual eFC predict clinical symptom

To further elucidate the relationship between extracted neural timescale and clinical symptoms, as well as to validate the robust biological significance of specific episode patterns, we utilized a multivariate approach to predict clinical measures based on these patterns. More specifically, to predict individual symptom severity, including HAMD and HAMA scores, we trained support vector regression (SVR) models using as input the neural timescales previously identified as significantly altered between patients and healthy controls (Liu et al., 2021). Functional features were partitioned into training and test sets using a 10-fold cross-validation strategy to mitigate overfitting. In each fold, a linear-kernel SVR model (LIBSVM, https://www.csie.ntu.edu.tw/~cjlin/libsvm/, default parameters) was fitted on the training data and applied to the held-out test set. Predictive performance was quantified as the Pearson correlation between predicted and observed symptom scores. To ensure robustness, the entire cross-validation procedure was repeated 1,000 times, and averaged predictions across iterations were used in final analyses. In parallel, regional feature importance contributing to symptom prediction was assessed using a random forest regression model.

## DATA AVAILABILITY

The clinical data could be accessed according to reasonable requests for corresponding authors. The raw fMRI data and MRI data for HCP was available on https://db.humanconnectome.org/. Heritability analyses were performed using Solar Eclipse 8.5.1b (https://www.solar-eclipse-genetics.org), Neuromap was available on (https://netneurolab.github.io/neuromaps/usage.html).

## CODE AVAILABILITY

Code will be available on https://github.com/Laoma29/Publication_codes.

## ACKNOWLEDGMENTS

Xiaobo Liu, Siyu Long, Jiadong Yan and Ke Xie are supported by the China Scholarship Council. Bin Wan is supported by International Max Planck Research School on Neuroscience of Communication: Function, Structure, and Plasticity (IMPRS NeuroCom), Graduate Academy Leipzig, and Mitacs Globalink Research Award. ZQL acknowledges support from the Fonds de Recherche du Qu\’ebec -- Nature et Technologies (FRQNT). Yujun Gao is supported by the Health of Hubei Province Scientific Research Project under Grant 2020Cfb512, and by the Mental Health Research Institute of Three Gorges University: YCXL-23-11.

## Supplementary Materials

### Research Participants Collection Procedures and Data Quality Control

A larger initial cohort underwent screening; however, 92 individuals were excluded based on predefined criteria to preserve data integrity and ensure methodological rigor. Within the BipD group, exclusions encompassed 9 participants who were subsequently reclassified with major depressive disorder (MDD) during longitudinal follow-up, 1 individual with an organic brain pathology, 2 participants unable to complete magnetic resonance imaging (MRI), 4 due to excessive head motion, and 4 owing to interference from metallic dental braces.

In the BipM cohort, 8 individuals were excluded due to motion artifacts, 2 due to diagnostic revision upon follow-up, 4 with comorbid gender identity disorder, 2 with underlying organic brain disease, and 8 who did not complete the MRI acquisition protocol. For the rBD group, exclusions included 2 participants who were later diagnosed with MDD and 6 due to motion-related artifacts exceeding acceptable thresholds.

In the healthy control (HC) group, 4 individuals were excluded due to the presence of dental braces, 4 due to excessive head motion (>2 mm) and refusal to undergo rescanning, and 15 due to a first-degree familial history of psychiatric or neurological disorders.

Following these exclusions, the final analytic sample comprised 42 individuals with BipD, 38 with BipM, 37 with rBD, 38 with mBD, and 35 healthy controls. All participants in the BipM cohort completed longitudinal assessments using multimodal methodologies (refer to Fig. S1 for procedural schema). This stringent screening and quality control framework underscores the critical importance of rigorous inclusion criteria and comprehensive data validation procedures to enhance the reliability, internal validity, and reproducibility of neuroimaging research in psychiatric populations.

---

**S-Figure.1.**
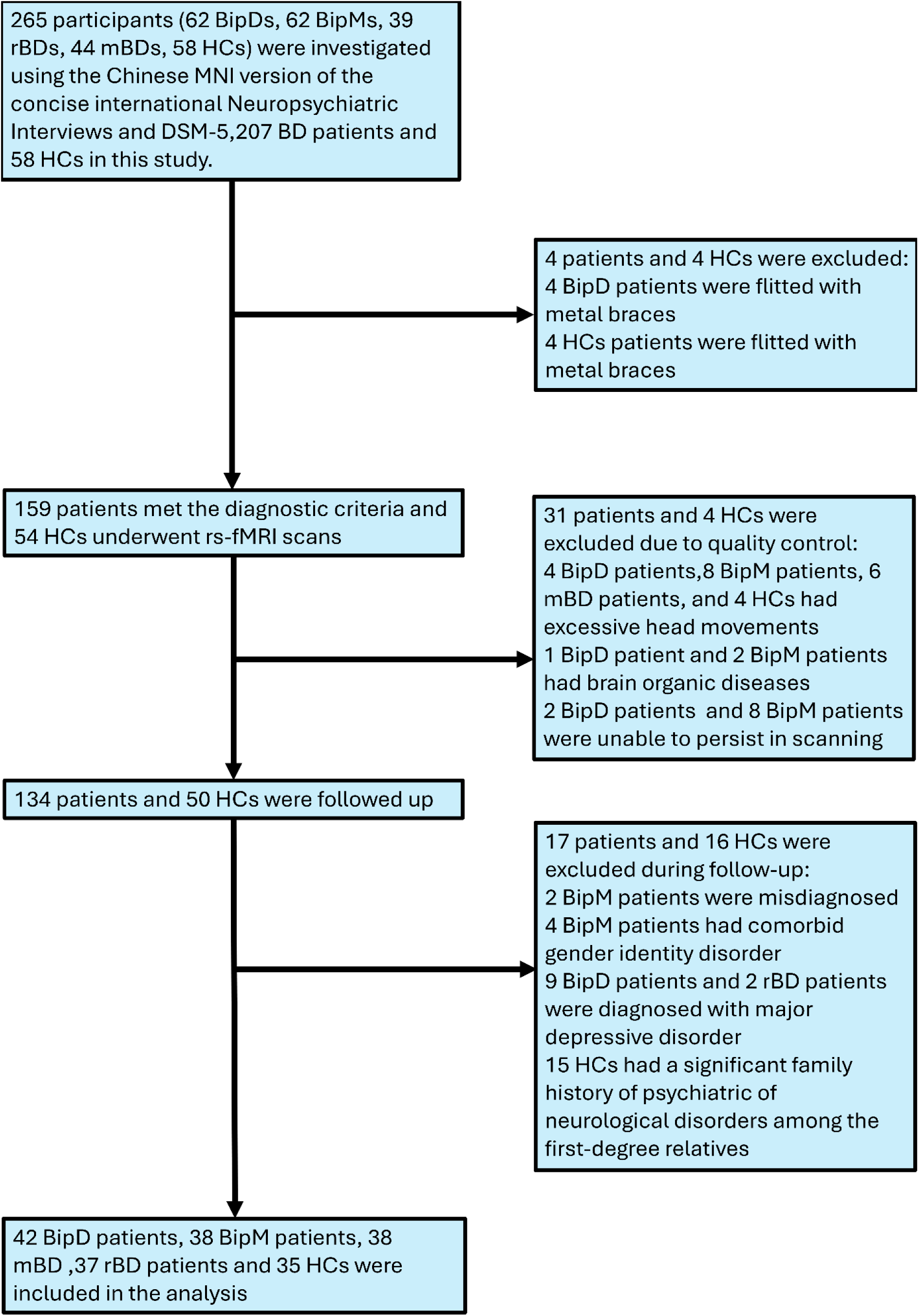
The research participants collection procedures and data quality control.

**S-Table1.** Demography (HDRS means Hamilton Depression Rating Scale, YMRS means Young Mania Rating Scale,PDQ means Perceived Disability Questionnaire, HAMA means Hamilton Anxiety Scale)

|  | *Age* | *Sex(M/F)* | *handnes*<br>*(R/L)* | *HDRS* | *YMRS* | *PDQ* | *HAMA* |
| --- | --- | --- | --- | --- | --- | --- | --- |
| *rBD* | 24.3±6.5 | 17/20 | 37/0 | 3.54±3.09 | 3.5±2.7 | 34.78±8.97 | 12.18±4.75 |
| *BipD* | 21.9±5.2 | 15/27 | 42/0 | 22.6±3.3 | 3.24±3.09 | 45.43±13.25 | 24.88±7.81 |
| *BipM* | 24.9±6.4 | 21/17 | 38/0 | 5.6±0.8 | 16.90±6.70 | 31.25±12.17 | 17.87±6.97 |
| *mBD* | 17.5±4.1 | 15/23 | 38/0 | 32.32±6.20 | 11.42±3.15 | ----- | ----- |
| *HC* | 26.7±7.3 | 14/21 | 35/0 | ----- | ----- | ----- | ----- |
| *F value* | 0.0307 | 0.0054 | -----<br>- | 321.05 | 111.61 | 15.67 | 35.67 |
| *p value* | 0.96 | 0.98 | -----<br>- | < 0.001 | < 0.001 | < 0.001 | < 0.001 |

